# Identification of transgene-free CRISPR edited plants of rice and tomato by monitoring DsRED fluorescence in dry seeds

**DOI:** 10.1101/533034

**Authors:** Norma Aliaga-Franco, Cunjin Zhang, Silvia Presa, Anjil K. Srivastava, Antonio Granell, David Alabadi, Ari Sadanandom, Miguel A. Blazquez, Eugenio G. Minguet

## Abstract

CRISPR/Cas technology for specific and precise modification of genomes has transformed molecular biology. However, quick elimination of the transgene remains a challenge in plant biotechnology after genome edition, especially for crops due to their long life cycle and multiploidy, not only to avoid transgene position effects and to minimize the probability of off-target mutation appearance, but also to deliver end users with edited plants free of the recombinant gene editing machinery. Counter selection based on resistance marker genes are inconvenient in the case of CRISPR/Cas applications because plants lacking the transgene cannot survive the selection, and thus two more generations must be screened to evaluate the presence of the transgene. In the case of some crops, generations can last between 6 months and a few years, and the workload may be a limiting factor because transgene detection by PCR requires germination of seeds, and selected plants must be grown until a new generation can be harvested. The expression of fluorescent proteins as selective markers has been successfully used in *Arabidopsis thaliana* (Stuitje et al., 2003) as a fast method for transgene presence detection prior to seed germination. Despite this clear advantage, it has not been tested in other species yet, because of the special requirements of *in vitro* transformation protocols. To overcome the above mentioned inconvenience, we have adapted fluorescence-dependent monitoring of transgene to genome editing approaches in tomato and rice with the goal of obtaining transgene-free homozygous edited crop plants in two generations.

Thanks to the modular design of construct generation through the Golden Braid cloning system (Vazquez-Vilar et al., 2016), we generated vectors containing the Cas9 transcriptional unit (TUs) under the control of optimal promoters for each species: AtUBQ10, ZmUBQ, and CaMV 35S for *Arabidopsis*, rice and tomato, respectively. Similarly, the sgRNA multiplexed TUs were placed under the control of the U6-26 promoter for Arabidopsis and tomato, and the U3 promoter for rice (Vazquez-Vilar et al., 2016). Specific resistance genes were also introduced into each vector for *in vitro* selection, as required by crop transformation protocols: hygromycin (Hyg) and kanamycin (Kan) for rice and tomato, respectively. Finally, an additional TU for expression of the fluorescent protein DsRED under the control of the At2S3 (Kroj et al., 2003) for Arabidopsis or CaMV 35S promoter for rice and tomato was added in each final vector (Fig. 1a).

**Figure 1.**
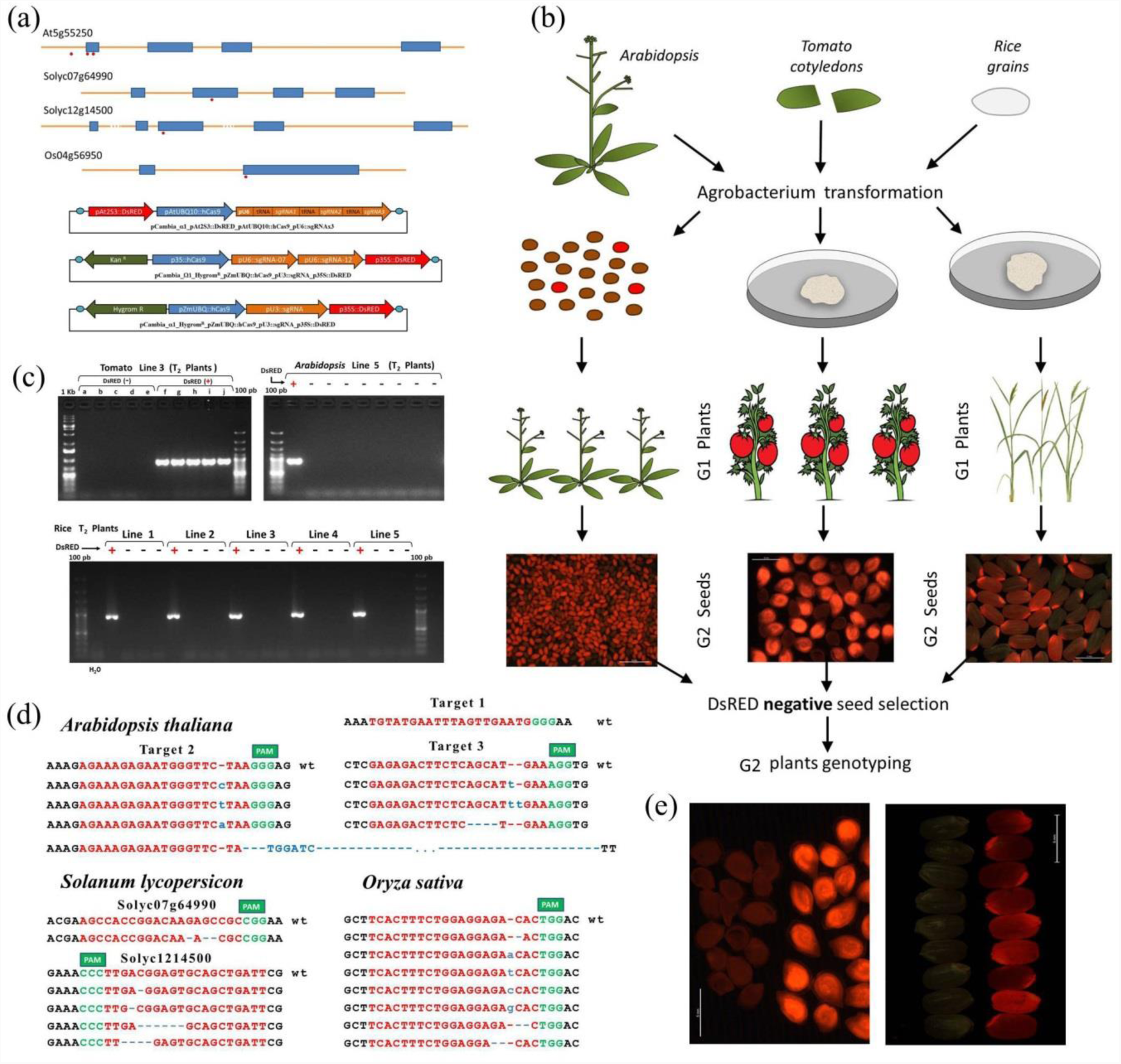
DsRED fluorescence as marker for transgene detection in dry seeds of tomato and rice. (a) Position of each guide RNA in the selected genes (At5g55250 [*Arabidopsis thaliana*], Solyc07g64990 and Solyc12g14500 [*Solanum lycopersicon*] and Os04g56950 [*Oryza sativa*]) and schemes of the Golden-Braid vectors generated for plant transformation. Transcriptional units are shown as arrows. (b) Diagram describing the steps for plant transformation: by floral dip in the case of Arabidopsis and in vitro transformation for tomato and rice. Selection of DsRED negative G2 seeds was done by direct visualization under a stereoscope equipped with DsRED filter. (c) PCR test confirming the absense of transgene in plants from DsRED-negative seeds, using Cas9 specific primers (5′-ggcggagcaagccaggaggaa-3′ and 5′-cttgacagccgcccccatcct-3′). Examples are shown for tomato, Arabidopsis and rice. (d) Mutations found in homozygosis in transgene-free G2 plants. CRISPR target sequence are shown in red, PAM sequence in green, and insertions in blue. (e) Ilustration of the identification of transgene-free tomato and rice seeds under white plus green light eith DsRED filter.

As a proof of concept, we decided to use the gene encoding IAA methyl transferase (IAMT) as an gene editing target in each species, given that loss of its function does not result in a differential visible phenotype to be selected for (Abbas et al., 2018). Phylogenetic analysis confirmed that *AtIAMT1* (At5g55250) had one ortholog in rice (Os04g56950) (Yang et al., 2008) and two in tomato (Solyc07g64990 and Solyc12g14500). Genomic DNA sequence for each selected gene in tomato and rice was analysed with ARES-GT tool (see below) for the identification and selection of one CRISPR guide RNA target for each gene. In the case of *Arabidopsis thaliana*, we selected three target sites of IAMT1 aiming to evaluate the efficiency of the vectors when looking for multiple mutations and deletions (Fig. 1a,d).

Several online tools are available for identification of CRISPR guide RNA targets, but cannot be used if the species of interest is not on the list of available genomes. Moreover, a list of selected targets based on a score threshold is usually provided which limits useful information. We have developed ARES-GT (https://github.com/eugomin/ARES-GT.git), a command line Python software (https://www.python.org/) that identifies all possible CRISPR targets (Cas9 or Cas12a/Cpf1) in DNA sequences and evaluates possible off-targets in a reference genome provided by user, thus non assembled contigs or not published genomes from any species can be used. ARES-GT algorithm selects the best sequences using two criteria: first, given that mismatches in the PAM proximal sequence (named seed sequence) have critical effect on DNA binding (Swarts and Jinek, 2018), any possible off-target with two or more mismatches in seed sequence is discarded; and second, the global number of mismatches is evaluated and any sequence above a user-defined threshold (user defined) is also discarded. ARES-GT generates a list of all CRISPR targets that only match its own sequence (the best candidates), but also a list with all the possible CRISPR targets with its possible off-targets locations with sequence alignment so, in the case of targeting a member of a gene family or with a very specific genome location, information will be available for evaluation and selection.

Each final construct (Fig. 1a) was introduced into the corresponding plant species following published transformation protocols: Arabidopsis was transformed by floral dip (Clough and Bent, 1998), whereas *in vitro* transformation protocols were used in rice (Nishimura et al., 2006) and tomato (Ellul et al., 2003) (Fig. 1b). As a result, 15 DsRED-positive T1 Arabidopsis seeds, 20 Hyg^R^ transgenic rice lines and 9 Kan^R^ transgenic tomato lines were selected for further analysis. Ideally, the ‘first generation’ of plants, that we call ‘G1’ for a uniform nomenclature, contains one copy of the transgene that will be segregated in the generation ‘G2’ independently of CRISPR induced germline mutations. While all G1 plants of rice and Arabidopsis produced seeds, two of the tomato plants were dwarf and did not produce any fruit and were discarded. Segregation analysis was done by visual observation of G2 dry seeds under a stereoscope equipped with DsRED filter and 12 Arabidopsis, 6 tomato, and 15 rice lines were retained for further analysis based on the 3:1 ratio of DsRED fluorescent vs non-fluorescent seeds. Plants from both types of seeds were grown under optimal greenhouse conditions until leaves were available for genomic DNA extraction. First, we confirmed that the Cas9 specific band was detected by PCR only in the plants from the DsRED positive seeds (Fig. 1c). Then each CRISPR-target region was PCR-amplified and sequenced from plants originated from non-fluorescent seeds. We confirmed the presence of mutations in all species and in all genes (Fig. 1d), indicating that stable mutations had been generated in the previous generation and inherited through the germline. In tomato, where two different genes were targeted at the same time, we identified different deletions in gene Solyc12g14500 in different plants both in heterozygosis and homozygosis, however we only detected a deletion of three nucleotides in Solyc07g64990 in one of the lines, which could be explained by deleterious effect of a null mutant but also because a low efficiency of sgRNA. In rice, all analysed lines did present mutations in the target gene, morever in a few of those lines all G2 plants presented the same mutation in homozygosis, indicating that the corresponding G1 plant likely was already homozygous. In *Arabidopsis*, mutations in two of the three targets were detected, including a 193-pb deletion between regions 2 and 3, most of them also in homozygosis in several plants. Those results are consistent with variable efficiency of sgRNAs reported in the literature, thus confirming that our vectors work as efficiently as other vector systems but with the advantage of using DsRED fluorescence as marker for transgene presence in dry seeds. It is noteworthy to mention that DsRED fluorescence have not decayed in seeds stored at least for one year.

In summary, we have demonstrated that the use of DsRED fluorescence as a selectable marker in seeds of crop plants facilitates the identification of transgene-free CRISPR/Cas9 edited plants in the second generation (Fig. 1e), minimizing the probability of off-target mutations. Golden Braid cloning system ensures easy modification of vectors for other Cas-like proteins (as Cas12a/Cpf1) or other fluorescence proteins. This approach can easily be extended to additional crops and model plants, as long as optimal promoters are available.

## Acknowledgements

We thank Diego Orzaez and Marta Vazquez for all the provided vectors from Golden Braid vector collection. Authors thank all the funding that has supported this work: NAF, DA, MAB and EGM thanks Spanish Ministry of Economy (AGL2014-57200-JIN, BFU2016-80621-P). SP and AG thanks Spanish Ministry of Economy (BIO2016-78601-R) and EC H2020 program, TRADITOM (634561), TomGEM (679796) Newcotiana (760331-2) and Pharma-Factory (SEP-210417525). CZ, AKS and AS thanks ERC project “SUMOrice” and BBSRC project “Flooding tolerance in rice”.

